# Impact of the sphingolipid metabolizing enzyme β-galactosylceramidase on mitochondrial sphingolipid profile and energetic metabolism in human melanoma cells

**DOI:** 10.64898/2026.08.18.745397

**Authors:** Davide Capoferri, Luca Mignani, Marzia Corli, Mirella Belleri, Anna Kovilakath, Lauren Ashley Cowart, Stefania Mitola, Marco Presta, Elisabetta Grillo

## Abstract

Mitochondrial plasticity, characterized by the dynamic balance between glycolysis and oxidative phosphorylation in response to genetic and microenvironmental changes, is a hallmark of melanoma progression. Sphingolipids play a significant role in various aspects of cancer cell biology, including metabolic reprogramming. Previous observations had shown that the lysosomal sphingolipid-metabolizing enzyme β-galactosylceramidase (GALC) rewires the lipid profile of mouse melanoma cells, exerting pro-oncogenic functions, gene silencing leading to a decreased oncogenic activity in murine and human melanoma cells. Here, we have focused on the mitochondrial sphingolipid composition and energetic metabolism in *GALC* knockout (KO) A2058 human melanoma cells. Targeted analysis of the mitochondrial sphingolipid profile, transcriptomic data, and mitochondrial structural and functional studies indicate that GALC loss drives a sphingolipid-mediated reprogramming of mitochondrial metabolism in absence of major structural alterations, characterized by bioenergetic insufficiency possibly due to ceramide- and sphingomyelin-driven impairment of respiratory chain function. Overall, these data indicate that *GALC* KO leads to a sphin-golipid-driven mitochondrial metabolic suppression and may provide novel information for the development of efficacious approaches in mitochondrial targeting melanoma therapies.

## 1. Introduction

Metastatic melanoma represents the deadliest form of skin cancer [1,2]. Numerous studies have demonstrated that metabolic reprogramming is a key driver of melanoma progression and metastasis, both of which are associated with profound alterations in cellular energy metabolism. Melanoma exhibits remarkable metabolic plasticity, dynamically shifting between glycolysis and oxidative phosphorylation in response to genetic and microenvironmental changes [3]. For example, the oncogenic BRAF^(V600E)^mutation, present in approximately 50% of human melanomas [4,5], promotes aerobic glycolysis while suppressing mitochondrial oxidative phosphorylation through activation of hypoxia-inducible factor 1α [6,7]. From a therapeutic standpoint, this metabolic flexibility may contribute to resistance to targeted therapies and immunotherapy, thereby limiting treatment efficacy in melanoma [8].

Within this framework, growing experimental evidence suggests that sphingolipids are key regulators of metabolic reprogramming through their effects on mitochondrial dynamics, cellular bioenergetics, apoptosis, and mitophagy [9]. Consequently, elucidating the impact of altered sphingolipid-metabolizing enzyme expression on mitochondrial plasticity and energetic metabolism may enhance our understanding of melanoma progression and support the development of innovative therapeutic strategies aimed at modulating mitochondrial dynamics [10].

The lysosomal acid hydrolase β-galactosylceramidase (GALC; EC 3.2.1.46) catalyzes the hydrolysis of β-galactose from β-galactosylceramide (GalCer) and other sphingolipids [11,12]. Evidence generated in our laboratory indicates that GALC may act as a pro-tumorigenic enzyme in human melanoma. Indeed, mRNA *in situ* hybridization analysis of human skin specimens revealed a progressive increase in *GALC* expression during the transition from common nevi to stage IV melanoma [13]. Accordingly, *Galc* downregulation impairs the tumorigenic and metastatic capacity of murine B16-F10 melanoma cells and is associated with marked alterations in their lipidomic profile. Likewise, *GALC* silencing reduces the tumorigenic potential of human melanoma cells [13]. Conversely, stable *GALC* overexpression induced by lentiviral transduction enhances the proliferative and migratory activity of *BRAF*-mutant A2058 and A375 human melanoma cells, accompanied by significant remodeling of the cell proteome [14,15]. In particular, transcriptomic and proteomic analyses indicated that *GALC* overexpression may exert a significant impact on mitochondrial plasticity in *BRAF*-mutant human melanoma cells [15].

Here, in the attempt to get further insights about the impact of GALC on the metabolic reprogramming of human melanoma, we have investigated the mitochondrial sphingolipid composition and energetic metabolism in *GALC* knockout (KO) BRAF^(V600E)^ mutated A2058 human melanoma cells. Analysis of the mitochondrial sphingolipid profile, transcriptomic data, and mitochondrial structural and functional studies converge on a model in which GALC loss drives a sphingolipid-mediated reprogramming of mitochondrial metabolism, characterized by bioenergetic insufficiency related to ceramide (Cer)- and sphingomyelin (SM)-driven impairment of respiratory chain function.

These findings shed new light on the tumorigenic role of GALC in human melanoma, identifying mitochondrial sphingolipid dysregulation as a previously unrecognized node in the pro-oncogenic cascade triggered by this sphingolipid metabolizing enzyme. Beyond their mechanistic significance, these results suggest that restoring mitochondrial sphingolipid homeostasis, or targeting the downstream bioenergetic vulnerabilities it creates, may represent a rational basis for the development of more effective mitochondria-directed therapeutic strategies in cancer.

## 2. Materials and Methods

### 2.1. Cell cultures and genome editing

A2058 cells were purchased from ATCC, grown in Dulbecco’s modified Eagle medium (DMEM; Thermo Fisher Scientific, Waltham, MA, USA) supplemented with 10% heat-inactivated fetal bovine serum (FBS), 100 U/mL penicillin, and 100 μg/mL streptomycin (Thermo Fisher Scientific) and maintained at 37 °C and 5% CO^2^ in a humidified incubator. Cells were regularly checked to exclude mycoplasma contamination. To generate *GALC* KO cells, The CRISPR/Cas9 approach was used targeting a 45-base pair-long sequence spanning between the *GALC* gene exon 7 and the 3’ intron using the following guides: GALC-c: AGGGTAAGGCATGTTTTACT and GALC-d: ATCAACCACCTTGAAGAGTT. Forty-eight hours upon transfection, A2058 cells were single-cell sorted by FACS Aria into 20% serum-enriched medium in 96-well plates, and allowed to grow for 4 weeks, replacing fresh culture medium every 4 or 5 days. Genomic PCR for the *GALC* locus performed with gGALC-fw: AATTATCAAGGTCTCCAGCGAG and gGALC-rev: TGAGAATGTAATCAAATGGGGA primers that amplify a 212-base pair-long amplicon 85 and 82 base pair distant from the pairing nucleotide of GALC-c and GALC-d guides, respectively. Control gL1CAM-fw: GGACTTTGATCTCATAGGGCAC and gL1CAM-rev: CAACCAACTCCTCTTCTGCTG primers were designed to amplify a 189-base pair-long amplicon for the locus of the *L1CAM* gene on chromosome X. In all the assays, control and KO cells were maintained in DMEM *plus* 2% FBS before starting the experimental procedure unless specified otherways.

### 2.2 GALC activity assay

GALC-mediated hydrolysis of the fluorescent GALC substrate LRh-6-GalCer (Nlissamine-rhodaminyl-6-aminohexanoylgalactosyl ceramide) following its incubation with 20 μg of cell extract was quantified by thin-layer chromatography (TLC) [16].

### 2.3. Cell proliferation assay

Cells were seeded in 48 well plates in DMEM supplemented with 2% FBS. After 24 hours, the medium was replaced with fresh cell culture medium containing 2% FBS and cells were counted at different time points thereafter.

### 2.4. Cell migration assay

Confluent cells maintained in DMEM supplemented with 2% FBS were scraped with a 200 μl tip to obtain a 2.0 mm thick denuded area. After 24 hours, wounded monolayers were photographed, and the width of the wounds was quantified by computerized analysis of the digitalized images in 8-10 independent sites per group [15].

### 2.5. Isolation of mitochondria for lipidomic analysis

Mitochondria were isolated using a modified protocol based on a previously described method [17]. Cells were scraped into 300 μl of ice-cold STM buffer [250 mM sucrose, 50 mM Tris-HCl, 5 mM MgCl_2_] and homogenized using a tight-fitting Teflon pestle in a Potter S homogenizer (Sartorius Stedium, Goettingen, Germany) at 1,000 rpm for 1 min. The homogenate was decanted into a pre-chilled centrifuge tube and maintained on ice for 30 min, with vortexing at maximum speed for 15 seconds every 2-3 min. The homogenate was centrifuged at 800 x g for 15 min at 4°C to remove nuclei and unbroken cells. The resulting supernatant was transferred to a new tube and centrifuged at 11,000 x g for 10 min at 4°C. The supernatant, representing the cytosolic fraction, was discarded, and the mitochondrial pellet was resuspended in 200 μl STM buffer and centrifuged at 11,000 x g for 10 min at 4°C. The final mitochondrial pellet was resuspended in 50 μL SOL buffer [50 mM Tris-HCl, 1 mM EDTA, 0.5% Triton-X-100] and subjected to brief sonication on ice at high setting for 10 seconds with 30 second pauses to lyse the mitochondrial membrane.

### 2.6. Targeted lipidomics

Lipids were extracted as described [18]. A 2:1:0.1 ratio of methanol:chloroform:water was added to equal amounts of starting homogenized cell or mitochondrial material, using bicinchoninic acid (BCA). In a screw cap 13 × 100 mm screw-top glass tube (VWR, 53283-800) containing 10 μl of internal standard solution [Avanti Polar Lipid standards were added as a cocktail containing 250 pmol each in 10 μl ethanol:methanol:water (7:2:1). Sphingoid base and sphingoid base 1-phosphate standards were 17-carbon chain-length analogs: C17-sphingosine, (2S,3R,4E)-2-aminoheptadec-4-ene-1,3-diol (d17:1-So); C17-sphingosine 1-phosphate, heptadecasphing-4-enine-1-phosphate (d17:1-So1P). Standards for N-acyl sphingolipids were C12-fatty acid analogs: C12-Cer, N-(dodecanoyl)-sphing-4-enine (d18:1/C12:0); C12-Cer 1-phosphate, N-(dodecanoyl)-sphing-4-enine-1-phosphate (d18:1/C12:0-Cer1P); C12-SM, N-(dodecanoyl)-sphing-4-enine-1-phosphocholine (d18:1/C12:0-SM); and C12-glucosylceramide, N-(dodecanoyl)-1-β-glucosyl-sphing-4-eine]. Tubes were sonicated for 30 seconds each, capped and incubated overnight in a water bath at 48°C. The tubes were centrifuged at 5000 × g for 10 min at 4°C, and the supernatant decanted into clean 13 × 100 mm borosilicate glass tubes, and the solvent evaporated under vacuum in a Speed-Vac (45°C). The dried lipids were then resuspended in 250 μl of LC-MS grade methanol by vortexing for 10 seconds, followed by immersing tubes in a sonicator water bath for 20–30 seconds. Insoluble debris was then removed by centrifugation at 5000 × g for 10 min at 4°C, and the supernatant was carefully decanted into an autoinjector vial (VWR, #46610-724), capped (VWR, #89239-020), and stored at −80°C until mass spectrometry analysis.

Analysis was performed as previously described [18,19]. For LC-ESI-MS/MS analyses, a Shimadzu Nexera LC-30 AD binary pump system coupled to a SIL-30AC autoinjector and DGU20A5R degasser coupled to an AB Sciex 5500 quadrupole/linear ion trap (QTrap) operating in a triple quadrupole mode were used. Q1 and Q3 were set to pass molecularly distinctive precursor and product ions (or a scan across multiple m/z in Q1 or Q3), using N2 to collisionally induce dissociations in Q2 (which was offset from Q1 by 30–120 eV). Sphingolipids were separated by reverse-phase LC on a Supelco 2.1 × 50 mm Ascentis Express C18 column (Millipore Sigma) at 35°C using a binary solvent system (flow rate of 0.5 mL/min). Prior to injection, the column was equilibrated for 30 seconds with a solvent mixture of 95% mobile phase A1 (CH3OH:water:HCOOH, 58:41:1, v:v:v, with 5 mM ammonium formate) and 5% mobile phase B1 (CH3OH:HCOOH, 99:1, v:v, with 5 mM ammonium formate), and after sample injection (typically 5 μl), the A1/B1 ratio was maintained at 95/5 for 2.25 minutes, followed by a linear gradient to 100% B1 over 1.5 minutes, which was held at 100% B1 for 5.5 minutes, followed by a 30 second gradient return to 95/5 A1/B1. The column was re-equilibrated with 95/5 A1/B1 for 30 seconds before the next run.

### 2.7. Mitochondria and Seahorse analyses

For mithocondrial analysis, cells seeded in 6-well plates (3 × 10^5^ cells/plate) were incubated in the dark for 20 min at 37°C with 200 nM MitoTracker Green FM, 5.0 μM MitoSOX™, 10 μM ThiolTracker™ Violet, or 1.0 μM TMRE All reagents from Invitrogen/ThermoFisher Scientific, USA). After washing, cells were suspended in medium and fluorescence detected by MACSQuant Analyzer (MiltenyiBiotec).

ATP levels in the cell extracts were evaluated with the ATP determination kit (A22066, Molecular probes, Invitrogen), according to manufacturer’s instructions.

The levels of mithocondrial OXPHOS complexes were assessed in total cell lysates by Western blotting probed with an anti-total OXPHOS antibody cocktail (ab110411, Abcam, Cambridge, UK).

Seahorse analysis: cells (4 × 10^4^ cells/well) were seeded on Seahorse XFe24 culture plates (Agilent, Santa Clara, CA, USA) the day before the assay in complete medium. The day of the assay, the medium was replaced with XF Base Medium (Agilent) *plus* 1 mM pyruvate/2 mM glutamine/10 mM glucose, pH 7.4. Oxygen consumption rate (OCR) measurements were performed at 6 min intervals (2 min mixing, 2 min waiting and 2 min measuring) using a Seahorse XFe24 Extracellular Flux Analyzer (XFeWave software) (Agilent, Santa Clara, CA, USA). Seahorse XF Mito-Stress Test was used to measure key parameters of mitochondrial function. Sequential treatments with oligomycin (1 μM final), FCCP (1 μM final), and rotenone/antimycin A (0.5 μM final) were performed to enable quantification of basal OCR, ATP-coupled OCR, maximal and spare respiration and proton leak. The data were normalized on total protein amounts measured using BCA assay.

### 2.8. Microarray Gene Expression analysis

Total RNA was obtained from three GALC KO and three control A2058 clones by using RNeasy mini kit (Qiagen Inc., Valencia, CA, USA) according to manufacturer instructions. Labelling of samples and hybridization to the Affymetrix Human Clariom S microarray chips (ThermoFisher Scientific, Waltham, MA, USA) covering over 20,000 transcripts were performed following the manufacturer’s protocols and data were obtained using the Affymetrix 5.0 software. Gene arrays were scaled to a median signal of 1500 and then analyzed independently. Signal calls were transformed to their base-2 logarithms, and each sample was normalized to have its mean and variance equal to the grand mean and average variance across all samples. The working dataset was generated upon SST-RMA summarization of the data in two groups by Transcriptome Analysis Console 4.0 (Applied biosystems).

## 3. Results

### 3.1. GALC knockout affects mithocondrial sphingolipid composition in human melanoma cells

To investigate the impact of GALC on the metabolic reprogramming in melanoma, CRISPR/Cas9 editing was performed to knockout the *GALC* gene in A2058 human melanoma cells. Single cell-derived control Cas9-null and KO cell clones (4 and 9 clones, respectively) were confirmed by genomic PCR (data not shown), RT-qPCR analysis of *GALC* mRNA levels, and enzymatic GALC activity assay (**Figure 1A**). As anticipated [13], experiments carried out on three independent *GALC* KO cell clones showed a decrease in cell proliferation and migration when compared to controls (**Figure 1B,C**), with no induction of cell apoptosis (**Figure 1D**).

**Figure 1.**
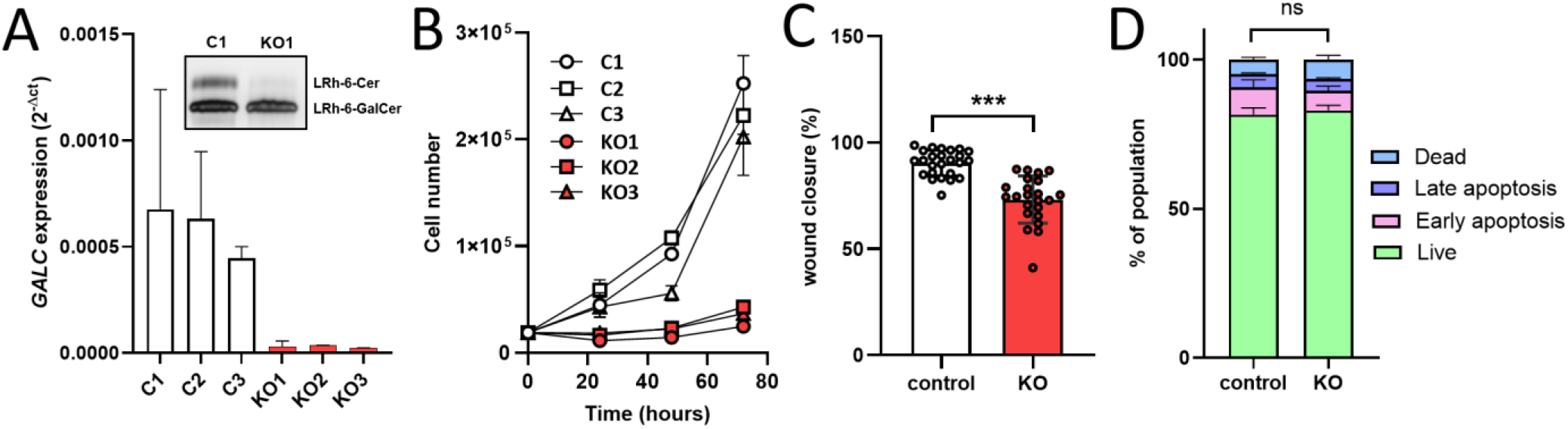
GALC KO in A2058 human melanoma cells. **A**) RT-qPCR analysis of GALC expression in three control (C1-C3) and three GALC KO (KO1-KO3) cell clones. Inset: GALC enzymatic TLC assay showing the incapacity of the KO1 cell extract to convert the LRh-6-GalCer substrate to the Lrh-6-Cer product when compared to control C1 cells. Similar results were obtained for all the clones tested. **B**) Cell proliferation of C1-C3 and KO1-KO3 cell clones grown under low serum conditions. **C**) Repair of mechanically wounded control and KO cell monolayers assessed 24 hours after wounding. **D**) Cell cycle analysis of control and KO cells. Data are the mean ± S.E.M (n = 3). In C and D, data from C1-C3 clones were pooled and compared with pooled data from KO1-KO3 clones. Student’s t test: ns, not significant; ^***^, P < 0.001.

*Galc* knockdown has been shown to exert a significant impact on the lipid profile of murine melanoma B16-F10 cells [13]. On this basis, randomly selected *GALC* KO and control cell clones (4 clones per group) were subjected to targeted sphingolipid analysis. The analysis included Cer, SM, monohexosylceramide (Hex-Cer), and lactosylceramide (LacCer) classes and their corresponding dihydro forms. Ten lipid species were analyzed for each class.

*GALC* KO did not exert a significant impact on the total levels of the sphingolipid classes analyzed, including dihydro lipids (data not shown). However, subtle changes emerged when individual sphingolipid Cer and SM species were examined: C22, C24, and C26:1 Cer species were significantly increased, with a concomitant reduction in C16 and C20 SM species. In contrast, no significant alterations were observed for any of the HexCer and LacCer species (**Figure 2**),

**Figure 2.**
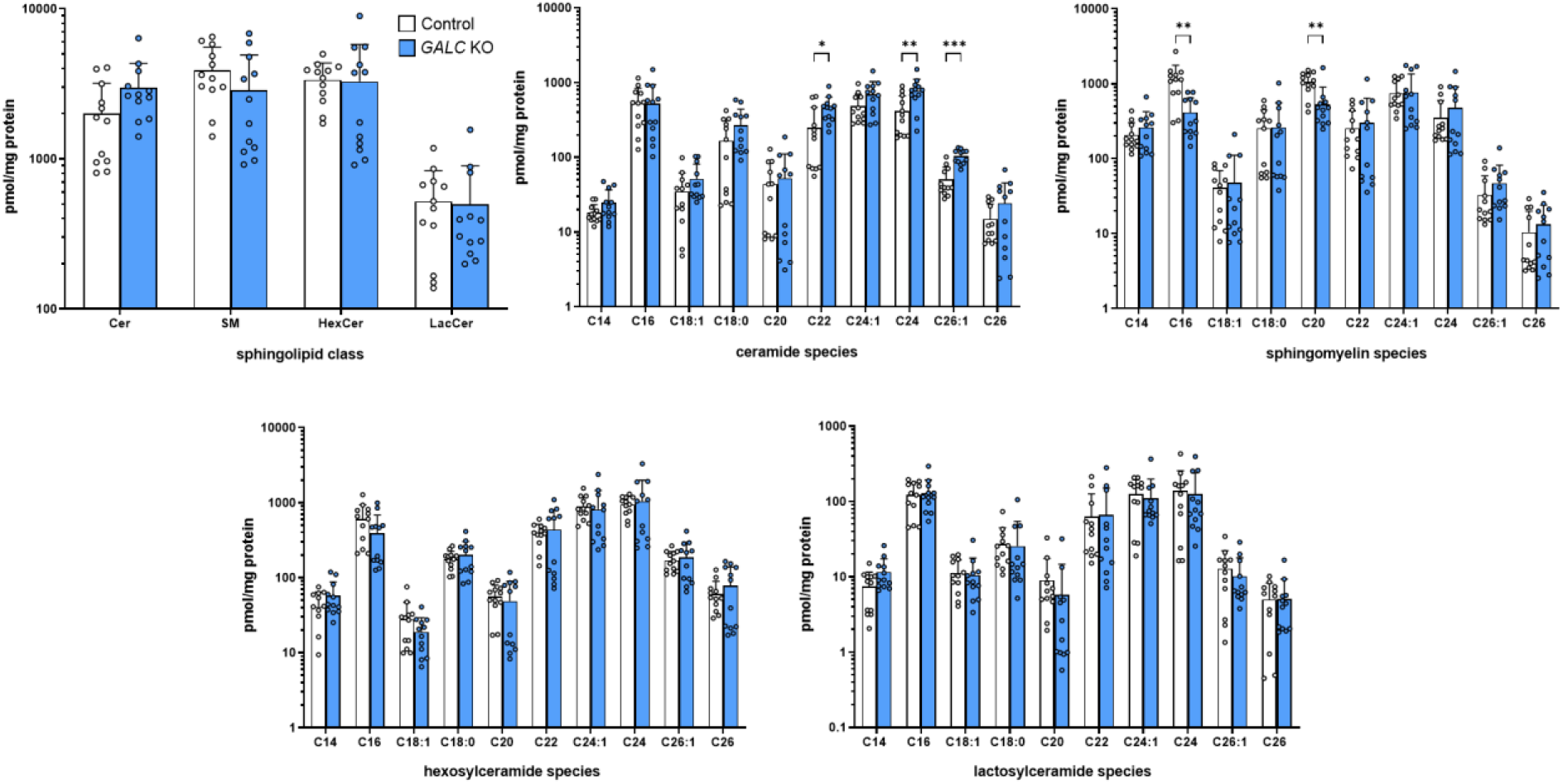
Impact of GALC KO on sphingolipid composition in melanoma cells. Targeted lipidomic analysis was performed in triplicate on control (white bars) and GALC KO (blue bars) cell clones (4 clones/group). Data are the mean ± S.E.M. Student’s t test: ^*^, P < 0.05; ^**^, P < 0.01; ^***^, P < 0.001.

We next applied lipidomic analysis to isolated mitochondria to investigate the impact of *GALC* KO on the mitochondrial sphingolipid composition. Mitochondria were isolated from *GALC* KO and control cell clones and subjected to evaluation of Cer, HexCer, LacCer, and SM content (**Figure 3)**.

**Figure 3.**
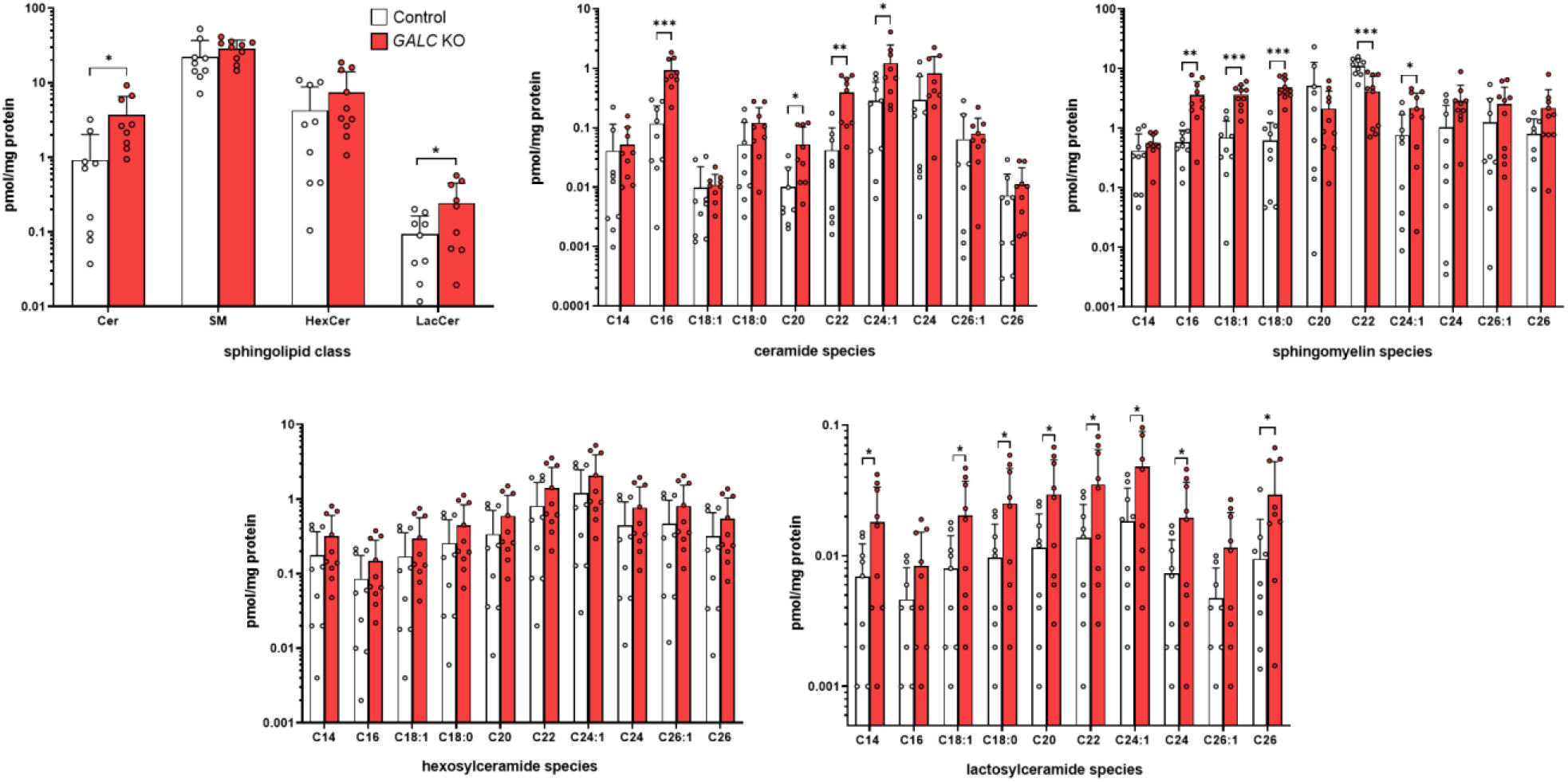
Impact of GALC KO on mitochondrial sphingolipid composition in melanoma cells. Targeted lipidomic analysis was performed in triplicate on mitochondria isolated from control (white bars) and GALC KO (red bars) cell clones (3 clones/group). Data are the mean ± S.E.M. Student’s t test: ^*^, P < 0.05; ^**^, P < 0.01; ^***^, P < 0.001.

*GALC* KO resulted in a significant increase in the mitochondrial levels of Cer and LacCer classes, whereas total SM and HexCer content remained unchanged. A more complex pattern emerged when individual sphingolipid species were examined. Specifically, C16, C20, C22, and C24:1 Cer species were significantly increased while C14, C24, C18:1, C18:0, and both C26 and C26:1 Cer species showed no significant changes. For SM, C16, C18:1, C18:0, C22, and C24:1 species were elevated, whereas all other species remained unchanged. In the case of LacCer, most species were increased, with the exceptions of C16 and C26:1 species, which were not significantly affected. By contrast, none of the HexCer species analyzed exhibited significant alterations. Overall, *GALC* KO promoted the accumulation of multiple Cer species and of the Cer-derived SM and LacCer species.

### 3.2. GALC KO affects gene-defined mitochondrial pathways in human melanoma cells

Next, transcriptomic analysis using an Affymetrix GeneChip array was performed on C1-C3 and KO1-KO3 cell clones. The analysis identified 360 upregulated and 509 downregulated genes when KO cell clones were compared to controls (**Figure 4A**). Gene expression data were used to interrogate the Mitocarta 3.0 database (https://www.broadinstitute.org/mitocarta) that represents an inventory of mammalian mitochondrial proteins and pathways in which gene sets are organized to define specific mitochondrial components, functions, and processes [20]. The analysis identified a list of terms defined by nuclear and/or mitochondrial genes that were significantly affected by *GALC* KO in A2058 cells (**Figure 4B**). When the differentially expressed gene clusters belonging to the pathways emerged from the Mitocarta 3.0 datasets were analyzed by the STRING tool (https://www.string-db.org), the most populated cluster (formed by 110 proteins, see **Supplementary Table 1**) was defined by enriched Gene Ontology (GO) terms that include, among others, the “Oxidative phosphorylation” and “Respiratory electron transport” terms (**Figure 4C**). The proteins from this cluster were then rendered by Pathview [21] on the “Oxidative phosphorylation” KEGG pathway (**Figure 4D**).

**Figure 4.**
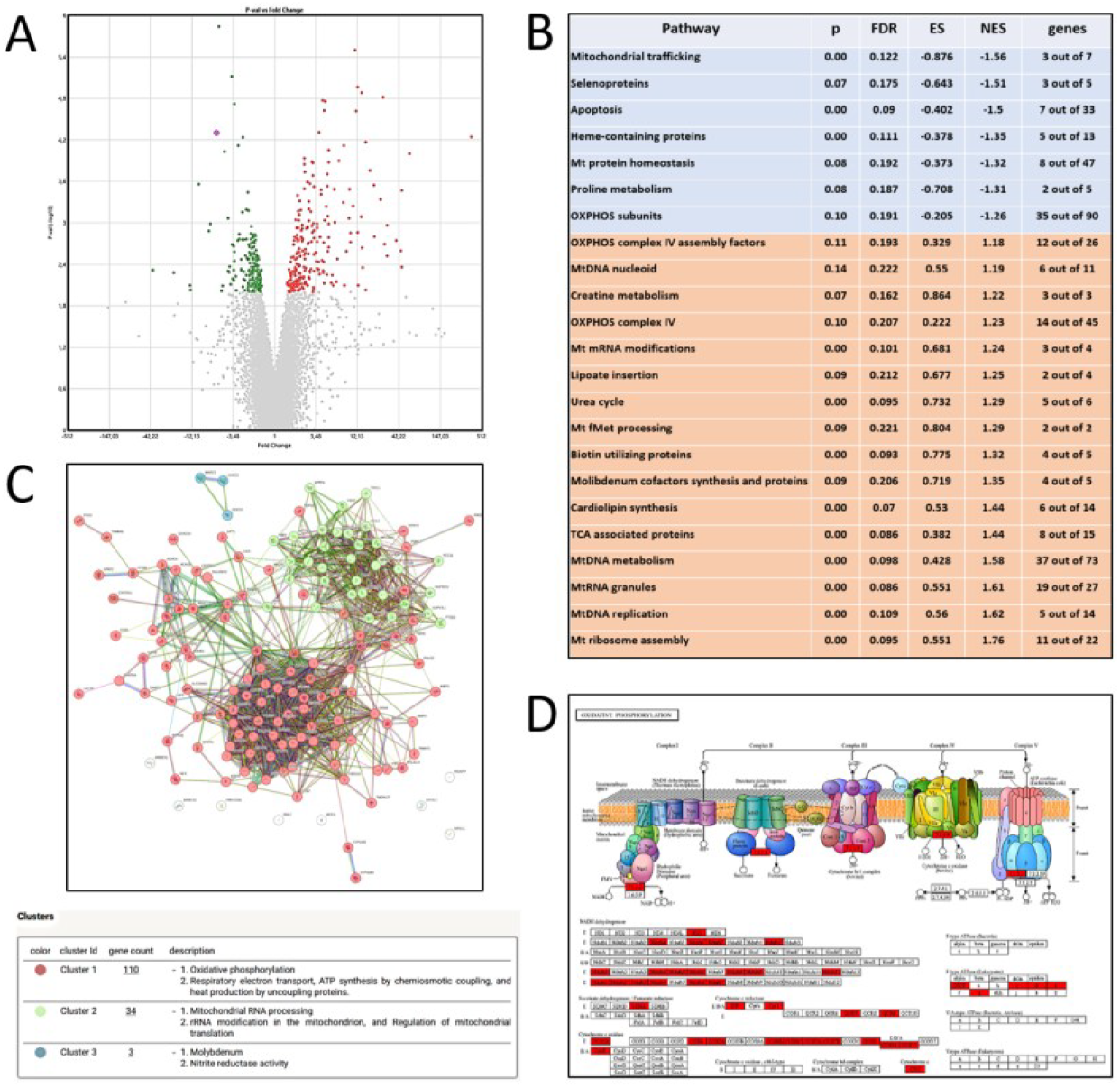
Mitochondrial pathways defined by the Mitocarta 3.0 database significantly affected by GALC KO in melanoma cells. **A**) Volcano plot analysis of the transcriptional profile of control and GALC KO cells. Upregulated and downregulated genes in GALC KO versus control cells are shown as red and green dots, respectively. **B**) Mitocarta-defined mitochondrial pathways significantly affected by GALC KO. **C**) STRING analysis of the differentially expressed genes in GALC KO versus control cells identified by analysis of the Mitocarta 3.0 database. **D**) Pathview rendering of the “Oxidative phosphorylation” KEGG pathway of the major gene cluster identified by STRING (in red in panel C).

### 3.3. GALC KO induces mitochondrial dysfunction in human melanoma cells

The effect of *GALC* KO on melanoma cell mitochondria at sphingolipid and transcriptional levels prompted us to investigate its impact on mitochondrial structure and functionality in human melanoma cells.

Mitochondria staining by the lipophilic cationic Mitotracker dye did not show any difference in the mitochondrial mass between *GALC* KO and control cells (**Figure 5A**). No significant difference was observed between control and *GALC* KO A2058 cells also when the mitochondrial subunits that compose the five complexes essential for the normal electron transfer chain were analyzed by Western blotting (**Figure 5B**). This was accompanied by a trend toward increased mitochondrial membrane depolarization following staining with the TMRE mitochondrial potential dye, although this effect did not achieve statistical significance. (**Figure 5C**). Nevertheless, *GALC* KO caused a dramatic decrease in ATP levels, paralleled by a decrease in GSH levels and reduced mitochondrial ROS production as assessed by staining with the MitoSOX probe (**Figure 5D-F**).

**Figure 5.**
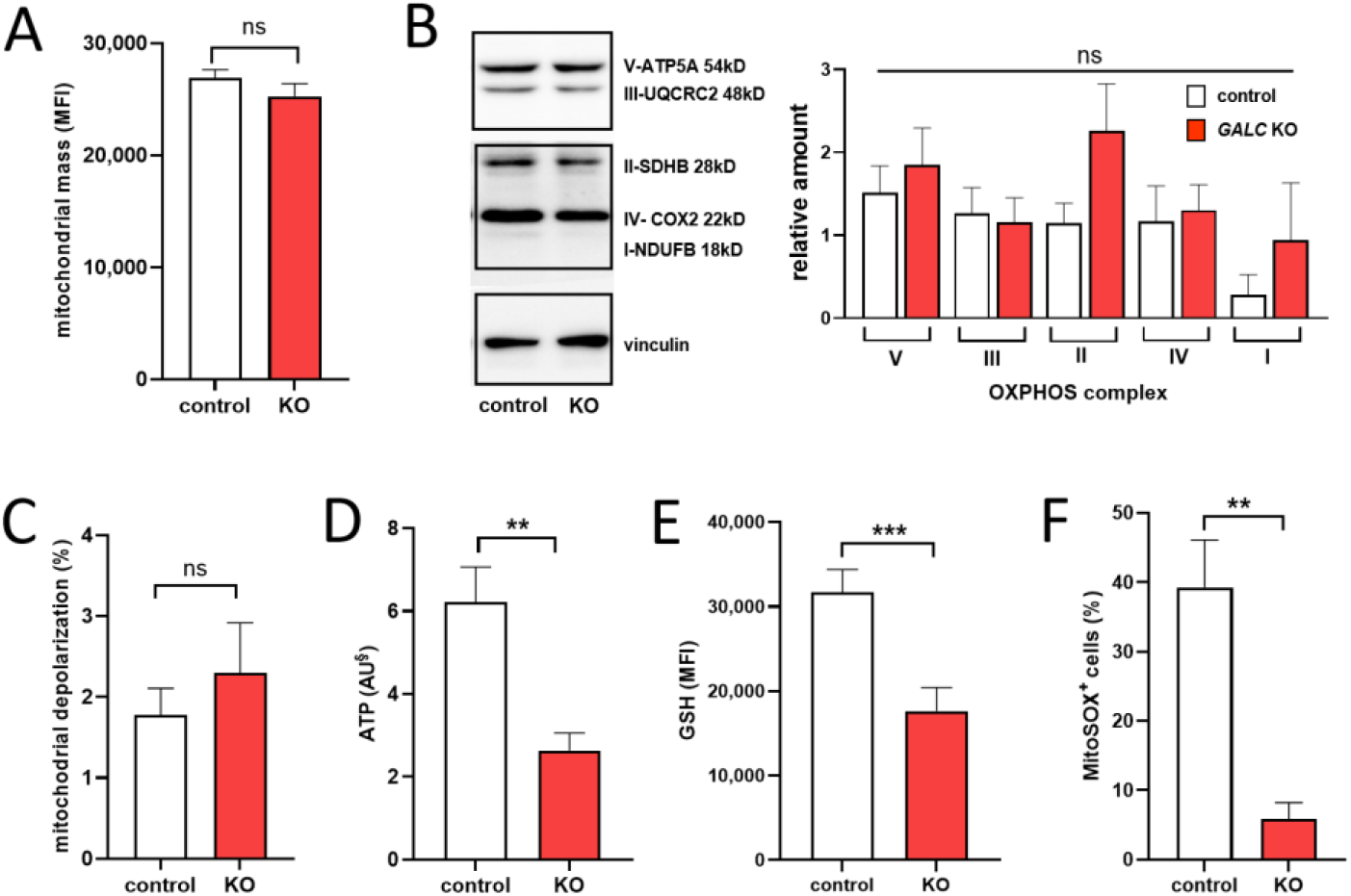
Impact of GALC KO on mitochondrial structure and functionality in melanoma cells. **A**) Evaluation of mitochondrial mass in control (white bars) and KO cells (red bars) by FACS analysis after staining with the Mitotracker dye. **B**) Western blot analysis of OXPHOS complexes in control and *GALC* KO cells. On the left, representative image of the results obtained with one control and one KO clone. Densitometric analysis of the Western blot data obtained with C1-C3 *versus* KO1-KO3 clones. **C**) Evaluation of mitochondrial membrane depolarization in control and *GALC* KO cells by FACS analysis after staining with the TMRE dye. **D**) Intracellular ATP levels in control and *GALC* KO cells. Evaluation of intracellular GSH levels (**E**) and mitochondrial ROS production (**F**) in control and KO cells by FACS analysis after staining with Thioltracker and MitoSOX probes, respectively. In all panels, data from C1-C3 clones were pooled and compared with the pooled data from KO1-KO3 clones. Data are the mean ± S.E.M. Student’s *t* test: ns, not significant; ^**^, P <0.01; ^***^, P < 0.001.

### 3.4. GALC KO affects oxidative metabolism and glycolysis in human melanoma cells

Given the impact of GALC KO on mitochondrial structure functionality, the Seahorse Mito-Stress Test was carried out on *GALC* KO *versus* control cells. To this end, nine *GALC* KO A2058 cell clones were pooled into three independent KO cell populations, each comprising three distinct clones, and compared to a pool of 3 control cell clones. To enable the characterization of key bioenergetic parameters, oxygen consumption rate (OCR) was measured under basal conditions and following the sequential injection of the ATP synthase inhibitor oligomycin, the uncoupling agent FCCP, and the complex I and III inhibitors rotenone/antimycin A. As shown in **Figure 6**, Consistent with the reduced ROS and ATP levels, *GALC* KO A2058 cells exhibited a marked reduction in mitochondrial respiration, as evidenced by decreased basal, proton leak, ATP-linked, and maximal OCR values (**Figure 6**).

**Figure 6.**
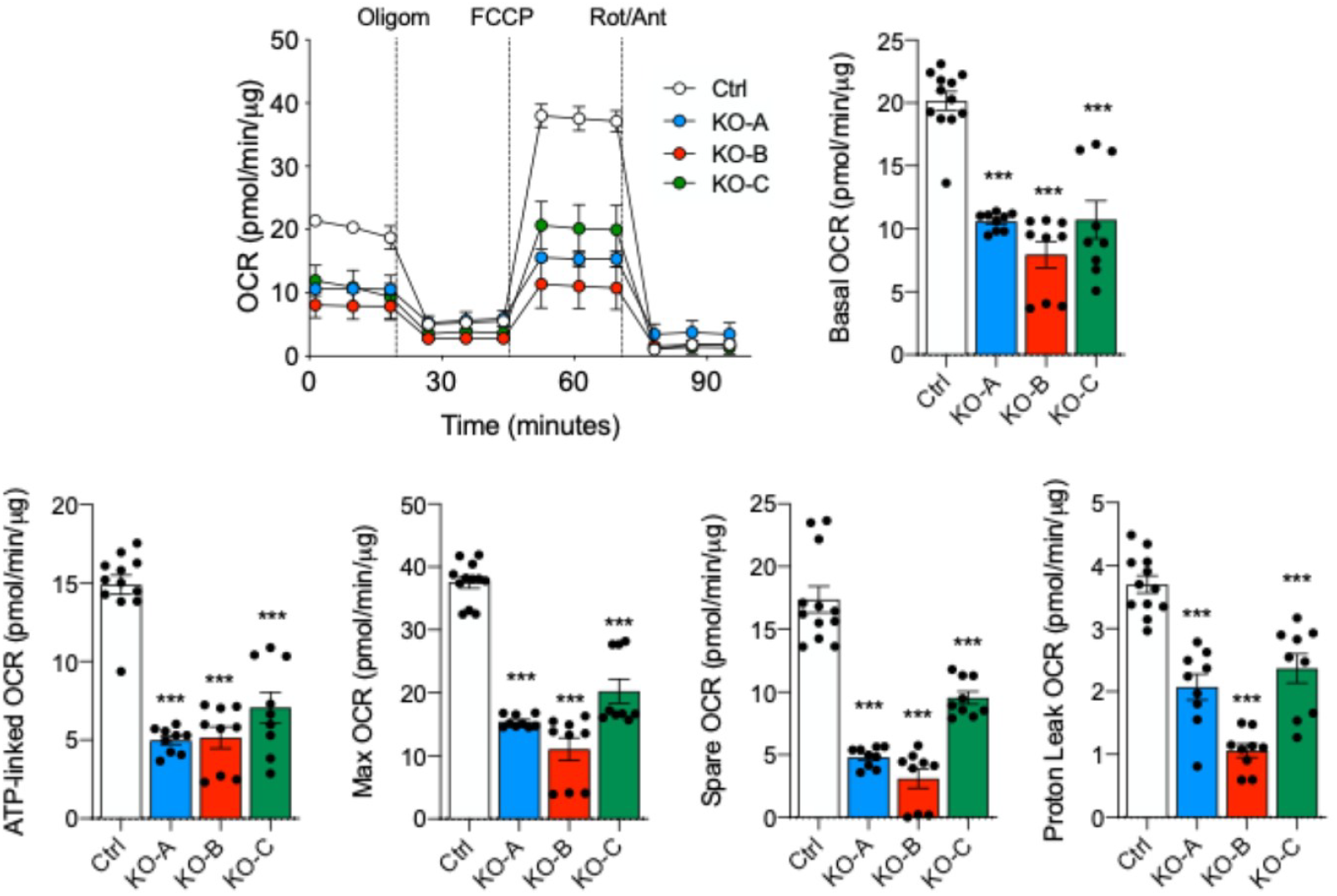
Impact of GALC KO on mitochondrial respiration in human melanoma cells. Seahorse analysis of cellular respiration (oxygen consumption rate, OCR) of *GALC* KO melanoma cells (3 pools of 3 clones each) compared to control cells (pool of 3 clones) before and after the injection of ATP synthase inhibitor oligomycin, the uncoupling agent FCCP, and the complex I and III inhibitors rotenone/antimycin A. Basal OCR, proton leak OCR, ATP-linked OCR, maximal OCR, and spare OCR were calculated and are shown in bar graphs. Data are the mean ± S.E.M. One-way ANOVA test: ^***^, P < 0.001.

## 4. Discussion

Alterations in sphingolipid metabolism, including changes affecting the tumor suppressor ceramide, have profound effects on melanoma progression [22–25]. GALC is a lysosomal sphingolipid-metabolizing enzyme that catalyzes the hydrolytic removal of galactose from β-GalcCer and other terminal β-Gal-containing sphingolipids [11,12].

Previous studies have shown that GALC may exert pro-oncogenic functions in human melanoma [13–15], with possible effects on mitochondrial plasticity [15]. Here, we extended these observations by investigating the impact of *GALC* KO on mitochondrial functionality and energetic metabolism in human melanoma A2058 cells. Consistent with previous observations in *GALC* downregulated human melanoma cells [13], *GALC* KO resulted in reduced cell proliferation and motility, without significantly affecting cell cycle progression and apoptosis. Moreover, *GALC* KO did not alter total mitochondrial mass or the levels of mitochondrial OXPHOS complexes and induced a modest increase in mitochondrial membrane depolarization. Despite the absence of major mitochondrial structural alterations, *GALC* KO cells exhibited reduced intracellular ATP levels accompanied by the concomitant decrease in GSH and mitochondrial ROS, consistent with a rewiring of cellular redox homeostasis and reduced oxidative metabolism. Accordingly, Seahorse analyses revealed decreased OCR values in *GALC* KO cell clones compared with control cells.

These alterations were accompanied by a marked remodeling of the mitochondrial sphingolipid profile, characterized by increased levels of multiple Cer, SM, and LacCer species. Notably, this occurred despite the absence of significant changes in total Cer, HexCer, LacCer, and SM levels in whole cell lysates, where the effect of *GALC* KO was limited to alterations in a few Cer and SM species. The observed mitochondrial sphingolipid imbalance likely underlies the mitochondrial dysfunction and bioenergetic failure in *GALC* KO cells. Mitochondrial Cer has been shown to directly inhibit electron transport chain activity by targeting multiple OXPHOS complexes [9,26], reducing electron flux, and consequently lowering both basal and maximal respiration, consistent with the decreased OCR and spare respiratory capacity observed here. Concurrently, Cer-induced pore formation in the outer mitochondrial membrane and Cer-mediated proton leak across the inner mitochondrial membrane [9] may account for the mild but measurable mitochondrial depolarization in GALC null cells. The resulting decrease in electron flux through the respiratory chain, rather than blockade of specific respiratory complexes, could also explain the paradoxical reduction in ROS generation that occurs in *GALC* KO cells despite impaired oxidative phosphorylation. Beyond Cer, the concomitant accumulation of SM species in the mitochondrial membrane may further impair respiratory function by increasing membrane rigidity and displacing cardiolipin from its native lipid domains [27]. This will destabilize OXPHOS supercomplex assembly and reducing coupling efficiency [28], a mechanism that would explain the functional impairment of the respiratory chain in the presence of unaltered complex abundance. Together, these data indicate that *GALC* KO leads to a sphingolipid-driven mitochondrial metabolic suppression.

Human melanoma A2058 cells harbor the tumor driving BRAF^(V600E)^ mutation which is present in approximately 50% of human melanomas [4,5] and represents a major target in melanoma therapy [29]. The BRAF^(V600E)^ mutation has been shown to suppresses mitochondrial oxidative phosphorylation and to drive aerobic glycolysis [6,7]. Our data indicate that *GALC* KO can exert a further impact on mitochondrial function and energetic metabolism in a *BRAF* mutated background. Mitochondrial plasticity has been implicated in resistance to targeted therapies in melanoma [8]. A better understanding of the role of sphingolipids and sphingolipid-metabolizing enzymes, including GALC, in mitochondrial dynamics and energetic metabolism may therefore facilitate the development of more effective mitochondria-targeted therapeutic strategies in cancer [10].

## Supplementary Materials

Table S1: List of the major Mitocarta-defined cluster of genes whose expression is modulated by *GALC* KO in A2058 cells.

## Author Contributions

Conceptualization: D.C., L.M., E.G, and M.P.; methodology: M.M. and L.A.C.; investigation: D.C., M.M., L.M and A.K.; data curation: D.C., L.M. and A.K.; statistical analysis: D.C. and L.M; writing—original draft preparation: M.P. and E.G.; writing—review and editing: M.P. and E.G.; supervision: M.P.; funding acquisition: M.M., M.P. and E.G. All authors have read and agreed to the published version of the manuscript.

## Institutional Review Board Statement

Not applicable.

## Data Availability Statement

The data presented in this study are available in the Supplementary Material.

## Abbreviations

The following abbreviations are used in this manuscript:

Cer: ceramide
HexCer: hexoxylceramide
LacCer: lactosylceramide
SM: sphingomyelin
GALC: β-galactosylceramidase
GO: Gene Ontology
OCR: oxygen consumption rate

