## Supplementary Table S1 for "Impact of the sphingolipid metabolizing enzyme β-galactosylceramidase on mitochondrial sphingolipid profile and energetic metabolism in human melanoma cells"

| | | Supplementary Table S1. List of proteins forming the major cluster identified by STRING analysis of the Mitocarta-defined mitochondrial pathways significantly affected by GALC KO in A2058 cells shown in red in Figure 4C. |
|---|---|---|
| **Protein name** | **Protein identifier** | **Protein description** |
| **ABCB7** | **9606.ENSP00000253577** | ATP-binding cassette sub-family B member 7, mitochondrial; Could be involved in the transport of heme from the mitochondria to the cytosol. Plays a central role in the maturation of cytosolic iron-sulfur (Fe/S) cluster-containing proteins. |
| **ACACA** | **9606.ENSP00000483300** | Acetyl-CoA carboxylase 1; Cytosolic enzyme that catalyzes the carboxylation of acetyl- CoA to malonyl-CoA, the first and rate-limiting step of de novo fatty acid biosynthesis. This is a 2 steps reaction starting with the ATP-dependent carboxylation of the biotin carried by the biotin carboxyl carrier (BCC) domain followed by the transfer of the carboxyl group from carboxylated biotin to acetyl-CoA. |
| **ACACB** | **9606.ENSP00000341044** | Acetyl-CoA carboxylase 2; Mitochondrial enzyme that catalyzes the carboxylation of acetyl-CoA to malonyl-CoA and plays a central role in fatty acid metabolism. Catalyzes a 2 steps reaction starting with the ATP-dependent carboxylation of the biotin carried by the biotin carboxyl carrier (BCC) domain followed by the transfer of the carboxyl group from carboxylated biotin to acetyl-CoA. Through the production of malonyl-CoA that allosterically inhibits carnitine palmitoyltransferase 1 at the mitochondria, negatively regulates fatty acid oxidation (By similarity). |
| **ACLY** | **9606.ENSP00000466259** | ATP-citrate synthase; Catalyzes the cleavage of citrate into oxaloacetate and acetyl-CoA, the latter serving as common substrate for de novo cholesterol and fatty acid synthesis. In the N-terminal section; belongs to the succinate/malate CoA ligase beta subunit family. |
| **ALDH18A1** | **9606.ENSP00000360268** | Delta-1-pyrroline-5-carboxylate synthase; Bifunctional enzyme that converts glutamate to glutamate 5- semialdehyde, an intermediate in the biosynthesis of proline, ornithine and arginine. |
| **ALKBH1** | **9606.ENSP00000216489** | Nucleic acid dioxygenase ALKBH1; Dioxygenase that acts as on nucleic acids, such as DNA and tRNA. Requires molecular oxygen, alpha-ketoglutarate and iron. A number of activities have been described for this dioxygenase, but recent results suggest that it mainly acts as on tRNAs and mediates their demethylation or oxidation depending on the context and subcellular compartment. Mainly acts as a tRNA demethylase by removing N(1)-methyladenine from various tRNAs, with a preference for N(1)-methyladenine at position 58 (m1A58) present on a stem loop structure of tRNAs. |
| **ANGEL2** | **9606.ENSP00000355929** | Angel homolog 2. |
| **ATP23** | **9606.ENSP00000300145** | Mitochondrial inner membrane protease ATP23 homolog; ATP23 metallopeptidase and ATP synthase assembly factor homolog. |
| **ATP5MC3** | **9606.ENSP00000284727** | ATP synthase F(0) complex subunit C3, mitochondrial; Mitochondrial membrane ATP synthase (F(1)F(0) ATP synthase or Complex V) produces ATP from ADP in the presence of a proton gradient across the membrane which is generated by electron transport complexes of the respiratory chain. F-type ATPases consist of two structural domains, F(1) - containing the extramembraneous catalytic core and F(0) - containing the membrane proton channel, linked together by a central stalk and a peripheral stalk. |
| **ATP5ME** | **9606.ENSP00000306003** | ATP synthase subunit e, mitochondrial, N-terminally processed; Mitochondrial membrane ATP synthase (F(1)F(0) ATP synthase or Complex V) produces ATP from ADP in the presence of a proton gradient across the membrane which is generated by electron transport complexes of the respiratory chain. F-type ATPases consist of two structural domains, F(1) - containing the extramembraneous catalytic core, and F(0) - containing the membrane proton channel, linked together by a central stalk and a peripheral stalk. |
| **ATP5MG** | **9606.ENSP00000300688** | ATP synthase subunit g, mitochondrial; Mitochondrial membrane ATP synthase (F(1)F(0) ATP synthase or Complex V) produces ATP from ADP in the presence of a proton gradient across the membrane which is generated by electron transport complexes of the respiratory chain. F-type ATPases consist of two structural domains, F(1) - containing the extramembraneous catalytic core, and F(0) - containing the membrane proton channel, linked together by a central stalk and a peripheral stalk. |
| **ATP5PD** | **9606.ENSP00000301587** | ATP synthase subunit d, mitochondrial; Mitochondrial membrane ATP synthase (F(1)F(0) ATP synthase or Complex V) produces ATP from ADP in the presence of a proton gradient across the membrane which is generated by electron transport complexes of the respiratory chain. F-type ATPases consist of two structural domains, F(1) - containing the extramembraneous catalytic core, and F(0) - containing the membrane proton channel, linked together by a central stalk and a peripheral stalk. |
| **ATP5PO** | **9606.ENSP00000290299** | ATP synthase subunit O, mitochondrial; Mitochondrial membrane ATP synthase (F(1)F(0) ATP synthase or Complex V) produces ATP from ADP in the presence of a proton gradient across the membrane which is generated by electron transport complexes of the respiratory chain. F-type ATPases consist of two structural domains, F(1) - containing the extramembraneous catalytic core and F(0) - containing the membrane proton channel, linked together by a central stalk and a peripheral stalk. |

| | | |
|---|---|---|
| **BAD** | **9606.ENSP00000378040** | Bcl2-associated agonist of cell death; Promotes cell death. Successfully competes for the binding to Bcl-X(L), Bcl-2 and Bcl-W, thereby affecting the level of heterodimerization of these proteins with BAX. Can reverse the death repressor activity of Bcl-X(L), but not that of Bcl-2 (By similarity). Appears to act as a link between growth factor receptor signaling and the apoptotic pathways. |
| **BCL2L1** | **9606.ENSP00000365230** | Bcl-2-like protein 1; Potent inhibitor of cell death. Inhibits activation of caspases. Appears to regulate cell death by blocking the voltage- dependent anion channel (VDAC) by binding to it and preventing the release of the caspase activator, CYC1, from the mitochondrial membrane. Also acts as a regulator of G2 checkpoint and progression to cytokinesis during mitosis. Isoform Bcl-X(S) promotes apoptosis. |
| **BCL2L10** | **9606.ENSP00000453562** | Bcl-2-like protein 10; Promotes cell survival. Suppresses apoptosis induced by BAX but not BAK. |
| **BNIP3** | **9606.ENSP00000357625** | BCL2/adenovirus E1B 19 kDa protein-interacting protein 3; Apoptosis-inducing protein that can overcome BCL2 suppression. May play a role in repartitioning calcium between the two major intracellular calcium stores in association with BCL2. Involved in mitochondrial quality control via its interaction with SPATA18/MIEAP: in response to mitochondrial damage, participates in mitochondrial protein catabolic process (also named MALM) leading to the degradation of damaged proteins inside mitochondria. Physical interaction of SPATA18/MIEAP, BNIP3 and BNIP3L/NIX at the mitochondrial outer membrane. |
| **CA5A** | **9606.ENSP00000498065** | Carbonic anhydrase 5A, mitochondrial; Reversible hydration of carbon dioxide. Low activity. |
| **CKMT1A** | **9606.ENSP00000406577** | Creatine kinase, mitochondrial 1A. |
| **CKMT2** | **9606.ENSP00000404203** | Creatine kinase S-type, mitochondrial; Reversibly catalyzes the transfer of phosphate between ATP and various phosphogens (e.g. creatine phosphate). Creatine kinase isoenzymes play a central role in energy transduction in tissues with large, fluctuating energy demands, such as skeletal muscle, heart, brain and spermatozoa; Belongs to the ATP:guanido phosphotransferase family. |
| **CMC1** | **9606.ENSP00000418348** | COX assembly mitochondrial protein homolog; Component of the MITRAC (mitochondrial translation regulation assembly intermediate of cytochrome c oxidase complex) complex, that regulates cytochrome c oxidase assembly. |
| **COA3** | **9606.ENSP00000354762** | Cytochrome c oxidase assembly factor 3 homolog, mitochondrial; Core component of the MITRAC (mitochondrial translation regulation assembly intermediate of cytochrome c oxidase complex) complex, that regulates cytochrome c oxidase assembly. MITRAC complexes regulate both translation of mitochondrial encoded components and assembly of nuclear-encoded components imported in mitochondrion. Required for efficient translation of MT-CO1 and mitochondrial respiratory chain complex IV assembly; Belongs to the COA3 family. |
| **COA5** | **9606.ENSP00000330730** | Cytochrome c oxidase assembly factor 5; Involved in an early step of the mitochondrial complex IV assembly process. |
| **COA7** | **9606.ENSP00000360593** | Cytochrome c oxidase assembly factor 7; Required for assembly of mitochondrial respiratory chain complex I and complex IV; Belongs to the hcp beta-lactamase family. |
| **COX10** | **9606.ENSP00000261643** | Protoheme IX farnesyltransferase, mitochondrial; Converts protoheme IX and farnesyl diphosphate to heme O. Belongs to the UbiA prenyltransferase family. |
| **COX11** | **9606.ENSP00000299335** | Cytochrome c oxidase assembly protein COX11, mitochondrial; Exerts its effect at some terminal stage of cytochrome c oxidase synthesis, probably by being involved in the insertion of the copper B into subunit I. |
| **COX14** | **9606.ENSP00000326052** | Cytochrome c oxidase assembly protein COX14; Core component of the MITRAC (mitochondrial translation regulation assembly intermediate of cytochrome c oxidase complex) complex, that regulates cytochrome c oxidase assembly. Requires for coordination of the early steps of cytochrome c oxidase assembly with the synthesis of MT-CO1. |
| **COX15** | **9606.ENSP00000016171** | Cytochrome c oxidase assembly protein COX15 homolog; May be involved in the biosynthesis of heme A. Belongs to the COX15/CtaA family. |
| **COX4I2** | **9606.ENSP00000365243** | Cytochrome c oxidase subunit 4 isoform 2, mitochondrial; Component of the cytochrome c oxidase, the last enzyme in the mitochondrial electron transport chain which drives oxidative phosphorylation. The respiratory chain contains 3 multisubunit complexes succinate dehydrogenase (complex II, CII), ubiquinol- cytochrome c oxidoreductase (cytochrome b-c1 complex, complex III, CIII) and cytochrome c oxidase (complex IV, CIV), that cooperate to transfer electrons derived from NADH and succinate to molecular oxygen, creating an electrochemical gradient over the inner membrane that drives transmembrane transport. |
| **COX5A** | **9606.ENSP00000317780** | Cytochrome c oxidase subunit 5A, mitochondrial; Component of the cytochrome c oxidase, the last enzyme in the mitochondrial electron transport chain which drives oxidative phosphorylation. The respiratory chain contains 3 multisubunit complexes succinate dehydrogenase (complex II, CII), ubiquinol- cytochrome c oxidoreductase (cytochrome b-c1 complex, complex III, CIII) and cytochrome c oxidase (complex IV, CIV), that cooperate to transfer electrons derived from NADH and succinate to molecular oxygen, creating an electrochemical gradient over the inner membrane that drives transmembrane transport |

| | | |
|---|---|---|
| **COX6B2** | **9606.ENSP00000467266** | Cytochrome c oxidase subunit 6B2; Component of the cytochrome c oxidase, the last enzyme in the mitochondrial electron transport chain which drives oxidative phosphorylation. The respiratory chain contains 3 multisubunit complexes succinate dehydrogenase (complex II, CII), ubiquinol- cytochrome c oxidoreductase (cytochrome b-c1 complex, complex III, CIII) and cytochrome c oxidase (complex IV, CIV), that cooperate to transfer electrons derived from NADH and succinate to molecular oxygen, creating an electrochemical gradient over the inner membrane that drives transmembrane transport. |
| **COX6C** | **9606.ENSP00000429707** | Cytochrome c oxidase subunit 6C; Component of the cytochrome c oxidase, the last enzyme in the mitochondrial electron transport chain which drives oxidative phosphorylation. The respiratory chain contains 3 multisubunit complexes succinate dehydrogenase (complex II, CII), ubiquinol- cytochrome c oxidoreductase (cytochrome b-c1 complex, complex III, CIII) and cytochrome c oxidase (complex IV, CIV), that cooperate to transfer electrons derived from NADH and succinate to molecular oxygen, creating an electrochemical gradient over the inner membrane that drives transmembrane transport. |
| **COX7A1** | **9606.ENSP00000292907** | Cytochrome c oxidase subunit 7A1, mitochondrial; Component of the cytochrome c oxidase, the last enzyme in the mitochondrial electron transport chain which drives oxidative phosphorylation. The respiratory chain contains 3 multisubunit complexes succinate dehydrogenase (complex II, CII), ubiquinol- cytochrome c oxidoreductase (cytochrome b-c1 complex, complex III, CIII) and cytochrome c oxidase (complex IV, CIV), that cooperate to transfer electrons derived from NADH and succinate to molecular oxygen, creating an electrochemical gradient over the inner membrane that drives transmembrane transport. |
| **COX7A2L** | **9606.ENSP00000367938** | Cytochrome c oxidase subunit 7A-related protein, mitochondrial; Involved in the regulation of oxidative phosphorylation and energy metabolism (By similarity). Necessary for the assembly of mitochondrial respiratory supercomplex (By similarity). Belongs to the cytochrome c oxidase VIIa family. |
| **COX7B** | **9606.ENSP00000497474** | Cytochrome c oxidase subunit 7B, mitochondrial; Component of the cytochrome c oxidase, the last enzyme in the mitochondrial electron transport chain which drives oxidative phosphorylation. The respiratory chain contains 3 multisubunit complexes succinate dehydrogenase (complex II, CII), ubiquinol- cytochrome c oxidoreductase (cytochrome b-c1 complex, complex III, CIII) and cytochrome c oxidase (complex IV, CIV), that cooperate to transfer electrons derived from NADH and succinate to molecular oxygen, creating an electrochemical gradient over the inner membrane that drives transmembrane transport. |
| **COX7B2** | **9606.ENSP00000379784** | Cytochrome c oxidase subunit 7B2, mitochondrial; Component of the cytochrome c oxidase, the last enzyme in the mitochondrial electron transport chain which drives oxidative phosphorylation. The respiratory chain contains 3 multisubunit complexes succinate dehydrogenase (complex II, CII), ubiquinol- cytochrome c oxidoreductase (cytochrome b-c1 complex, complex III, CIII) and cytochrome c oxidase (complex IV, CIV), that cooperate to transfer electrons derived from NADH and succinate to molecular oxygen, creating an electrochemical gradient over the inner membrane that drives transmembrane transport. |
| **COX8C** | **9606.ENSP00000340568** | Cytochrome c oxidase subunit 8C, mitochondrial; Component of the cytochrome c oxidase, the last enzyme in the mitochondrial electron transport chain which drives oxidative phosphorylation. The respiratory chain contains 3 multisubunit complexes succinate dehydrogenase (complex II, CII), ubiquinol- cytochrome c oxidoreductase (cytochrome b-c1 complex, complex III, CIII) and cytochrome c oxidase (complex IV, CIV), that cooperate to transfer electrons derived from NADH and succinate to molecular oxygen, creating an electrochemical gradient over the inner membrane that that drives transmembrane transport. |
| **CPS1** | **9606.ENSP00000402608** | Carbamoyl-phosphate synthase [ammonia], mitochondrial; Involved in the urea cycle of ureotelic animals where the enzyme plays an important role in removing excess ammonia from the cell. |
| **CYC1** | **9606.ENSP00000317159** | Cytochrome c1, heme protein, mitochondrial; Component of the ubiquinol-cytochrome c oxidoreductase, a multisubunit transmembrane complex that is part of the mitochondrial electron transport chain which drives oxidative phosphorylation. The respiratory chain contains 3 multisubunit complexes succinate dehydrogenase (complex II, CII), ubiquinol-cytochrome c oxidoreductase (cytochrome b-c1 complex, complex III, CIII) and cytochrome c oxidase (complex IV, CIV), that cooperate to transfer electrons derived from NADH and succinate to molecular oxygen. |
| **CYCS** | **9606.ENSP00000307786** | Cytochrome c; Electron carrier protein. The oxidized form of the cytochrome c heme group can accept an electron from the heme group of the cytochrome c1 subunit of cytochrome reductase. Cytochrome c then transfers this electron to the cytochrome oxidase complex, the final protein carrier in the mitochondrial electron-transport chain. |
| **CYP11B1** | **9606.ENSP00000292427** | Cytochrome P450 11B1, mitochondrial; A cytochrome P450 monooxygenase involved in the biosynthesis of adrenal corticoids. Catalyzes the hydroxylation of carbon hydrogen bond at 11-beta position of 11-deoxycortisol and 11- deoxycorticosterone/21-hydroxyprogesterone yielding cortisol or corticosterone, respectively. Mechanistically, uses molecular oxygen inserting one oxygen atom into a substrate and reducing the second into a water molecule. |
| **CYP11B2** | **9606.ENSP00000325822** | Cytochrome P450 11B2, mitochondrial; A cytochrome P450 monooxygenase that catalyzes the biosynthesis of adrenal mineralocorticoid aldosterone. Catalyzes three sequential oxidative reactions of 11-deoxycorticosterone/21- hydroxyprogesterone, namely 11-beta hydroxylation followed with two successive oxidations at C18 to yield 18-hydroxy and then 18-aldehyde derivatives, resulting in the formation of aldosterone. Mechanistically, uses molecular oxygen inserting one oxygen atom into a substrate and reducing the second into a water molecule. |

| | | |
|---|---|---|
| **CYP27A1** | **9606.ENSP00000258415** | Sterol 26-hydroxylase, mitochondrial; Cytochrome P450 monooxygenase that catalyzes regio- and stereospecific hydroxylation of cholesterol and its derivatives. Hydroxylates (with R stereochemistry) the terminal methyl group of cholesterol side-chain in a three step reaction to yield at first a C26 alcohol, then a C26 aldehyde and finally a C26 acid. Regulates cholesterol homeostasis by catalyzing the conversion of excess cholesterol to bile acids via both the 'neutral' (classic) and the 'acid' (alternative) pathways. May also regulate cholesterol homeostasis. |
| **D2HGDH** | **9606.ENSP00000315351** | D-2-hydroxyglutarate dehydrogenase, mitochondrial; Catalyzes the oxidation of D-2-hydroxyglutarate to alpha- ketoglutarate. |
| **FAHD1** | **9606.ENSP00000372112** | Acylpyruvase FAHD1, mitochondrial; Probable mitochondrial acylpyruvase which is able to hydrolyze acetylpyruvate and fumarylpyruvate in vitro. Also has oxaloacetate decarboxylase activity. |
| **GATM** | **9606.ENSP00000379895** | Glycine amidinotransferase, mitochondrial; Catalyzes the biosynthesis of guanidinoacetate, the immediate precursor of creatine. Creatine plays a vital role in energy metabolism in muscle tissues. May play a role in embryonic and central nervous system development. May be involved in the response to heart failure by elevating local creatine synthesis; Belongs to the amidinotransferase family. |
| **GPAM** | **9606.ENSP00000265276** | Glycerol-3-phosphate acyltransferase 1, mitochondrial; Esterifies acyl-group from acyl-ACP to the sn-1 position of glycerol-3-phosphate, an essential step in glycerolipid biosynthesis. |
| **GPAT2** | **9606.ENSP00000389395** | Glycerol-3-phosphate acyltransferase 2, mitochondrial; Esterifies acyl-group from acyl-ACP to the sn-1 position of glycerol-3-phosphate, an essential step in glycerolipid biosynthesis. Required for primary processing step during piRNA biosynthesis. Molecular mechanisms by which it promotes piRNA biosynthesis are unclear and do not involve its acyltransferase activity. Belongs to the GPAT/DAPAT family. |
| **GPX4** | **9606.ENSP00000346103** | Phospholipid hydroperoxide glutathione peroxidase; Essential antioxidant peroxidase that directly reduces phospholipid hydroperoxide even if they are incorporated in membranes and lipoproteins (By similarity). Can also reduce fatty acid hydroperoxide, cholesterol hydroperoxide and thymine hydroperoxide (By similarity). Plays a key role in protecting cells from oxidative damage by preventing membrane lipid peroxidation (By similarity). Required to prevent cells from ferroptosis, a non-apoptotic cell death resulting from an iron-dependent accumulation of lipid reactive oxygen species. |
| **HIGD1A** | **9606.ENSP00000398064** | HIG1 domain family member 1A, mitochondrial; Proposed subunit of cytochrome c oxidase (COX, complex IV), which is the terminal component of the mitochondrial respiratory chain that catalyzes the reduction of oxygen to water. May play a role in the assembly of respiratory supercomplexes. |
| **IMMP2L** | **9606.ENSP00000384966** | Mitochondrial inner membrane protease subunit 2; Catalyzes the removal of transit peptides required for the targeting of proteins from the mitochondrial matrix, across the inner membrane, into the inter-membrane space. Known to process the nuclear encoded protein DIABLO; Belongs to the peptidase S26 family. IMP2 subfamily. |
| **L2HGDH** | **9606.ENSP00000267436** | L-2-hydroxyglutarate dehydrogenase, mitochondrial; L-2-hydroxyglutarate dehydrogenase; Belongs to the L2HGDH family. |
| **LACTB** | **9606.ENSP00000261893** | Serine beta-lactamase-like protein LACTB, mitochondrial; Mitochondrial serine protease that acts as a regulator of mitochondrial lipid metabolism. Acts by decreasing protein levels of PISD, a mitochondrial enzyme that converts phosphatidylserine (PtdSer) to phosphatidylethanolamine (PtdEtn), thereby affecting mitochondrial lipid metabolism. It is unclear whether it acts directly by mediating proteolysis of PISD or by mediating proteolysis of another lipid metabolism protein. Acts as a tumor suppressor that has the ability to inhibit proliferation of multiple types of breast cancer cells. |
| **LAP3** | **9606.ENSP00000226299** | Cytosol aminopeptidase; Presumably involved in the processing and regular turnover of intracellular proteins. Catalyzes the removal of unsubstituted N- terminal amino acids from various peptides; Belongs to the peptidase M17 family. |
| **LIAS** | **9606.ENSP00000492260** | Lipoyl synthase, mitochondrial; Catalyzes the radical-mediated insertion of two sulfur atoms into the C-6 and C-8 positions of the octanoyl moiety bound to the lipoyl domains of lipoate-dependent enzymes, thereby converting the octanoylated domains into lipoylated derivatives. |
| **LIPT1** | **9606.ENSP00000377115** | Lipoyltransferase 1, mitochondrial; Catalyzes the transfer of the lipoyl group from lipoyl-AMP to the specific lysine residue of lipoyl domains of lipoate-dependent enzymes; Belongs to the LplA family. |
| **LRPPRC** | **9606.ENSP00000260665** | Leucine-rich PPR motif-containing protein, mitochondrial; May play a role in RNA metabolism in both nuclei and mitochondria. In the nucleus binds to HNRPA1-associated poly(A) mRNAs and is part of nmRNP complexes at late stages of mRNA maturation which are possibly associated with nuclear mRNA export. May bind mature mRNA in the nucleus outer membrane. In mitochondria binds to poly(A) mRNA. Plays a role in translation or stability of mitochondrially encoded cytochrome c oxidase (COX) subunits. May be involved in transcription regulation. |
| **MCCC1** | **9606.ENSP00000265594** | Methylcrotonoyl-CoA carboxylase subunit alpha, mitochondrial; Biotin-attachment subunit of the 3-methylcrotonyl-CoA carboxylase, an enzyme that catalyzes the conversion of 3-methylcrotonyl-CoA to 3-methylglutaconyl-CoA, a critical step for leucine and isovaleric acid catabolism. |
| **ME2** | **9606.ENSP00000321070** | NAD-dependent malic enzyme, mitochondrial; Malic enzyme 2; Belongs to the malic enzyme family. |

| | | |
|---|---|---|
| **METAP1D** | **9606.ENSP00000315152** | Methionine aminopeptidase 1D, mitochondrial; Removes the N-terminal methionine from nascent proteins. The N-terminal methionine is often cleaved when the second residue in the primary sequence is small and uncharged (Met-Ala-, Cys, Gly, Pro, Ser, Thr, or Val). Requires deformylation of the N(alpha)-formylated initiator methionine before it can be hydrolyzed (By similarity). May play a role in colon tumorigenesis; Belongs to the peptidase M24A family. Methionine aminopeptidase type 1 subfamily. |
| **MIEF1** | **9606.ENSP00000385110** | Mitochondrial dynamics protein MID51; Mitochondrial outer membrane protein which regulates mitochondrial fission. Promotes the recruitment and association of the fission mediator dynamin-related protein 1 (DNM1L) to the mitochondrial surface independently of the mitochondrial fission FIS1 and MFF proteins. Regulates DNM1L GTPase activity and DNM1L oligomerization. Binds ADP and can also bind GDP, although with lower affinity. Does not bind CDP, UDP, ATP, AMP or GTP. Inhibits DNM1L GTPase activity in the absence of bound ADP. |
| **MT-ND5** | **9606.ENSP00000354813** | NADH-ubiquinone oxidoreductase chain 5; Core subunit of the mitochondrial membrane respiratory chain NADH dehydrogenase (Complex I) that is believed to belong to the minimal assembly required for catalysis. Complex I functions in the transfer of electrons from NADH to the respiratory chain. The immediate electron acceptor for the enzyme is believed to be ubiquinone (By similarity). |
| **MTPAP** | **9606.ENSP00000263063** | Poly(A) RNA polymerase, mitochondrial; Polymerase that creates the 3' poly(A) tail of mitochondrial transcripts. Can use all four nucleotides, but has higher activity with ATP and UTP (in vitro). Plays a role in replication-dependent histone mRNA degradation. May be involved in the terminal uridylation of mature histone mRNAs before their degradation is initiated. Might be responsible for the creation of some UAA stop codons which are not encoded in mtDNA. |
| **NAGS** | **9606.ENSP00000293404** | N-acetylglutamate synthase conserved domain form; Plays a role in the regulation of ureagenesis by producing the essential cofactor N-acetylglutamate (NAG), thus modulating carbamoylphosphate synthase I (CPS1) activity. Belongs to the acetyltransferase family. |
| **NDUFA1** | **9606.ENSP00000360492** | NADH dehydrogenase [ubiquinone] 1 alpha subcomplex subunit 1; Accessory subunit of the mitochondrial membrane respiratory chain NADH dehydrogenase (Complex I), that is believed not to be involved in catalysis. Complex I functions in the transfer of electrons from NADH to the respiratory chain. The immediate electron acceptor for the enzyme is believed to be ubiquinone. |
| **NDUFA12** | **9606.ENSP00000330737** | NADH dehydrogenase [ubiquinone] 1 alpha subcomplex subunit 12; Accessory subunit of the mitochondrial membrane respiratory chain NADH dehydrogenase (Complex I), that is believed not to be involved in catalysis. Complex I functions in the transfer of electrons from NADH to the respiratory chain. The immediate electron acceptor for the enzyme is believed to be ubiquinone. |
| **NDUFA4** | **9606.ENSP00000339720** | Cytochrome c oxidase subunit NDUFA4; Component of the cytochrome c oxidase, the last enzyme in the mitochondrial electron transport chain which drives oxidative phosphorylation. The respiratory chain contains 3 multisubunit complexes succinate dehydrogenase (complex II, CII), ubiquinol- cytochrome c oxidoreductase (cytochrome b-c1 complex, complex III, CIII) and cytochrome c oxidase (complex IV, CIV), that cooperate to transfer electrons derived from NADH and succinate to molecular oxygen, creating an electrochemical gradient over the inner membrane that drives transmembrane transport. |
| **NDUFA5** | **9606.ENSP00000417142** | NADH dehydrogenase [ubiquinone] 1 alpha subcomplex subunit 5; Accessory subunit of the mitochondrial membrane respiratory chain NADH dehydrogenase (Complex I), that is believed not to be involved in catalysis. Complex I functions in the transfer of electrons from NADH to the respiratory chain. The immediate electron acceptor for the enzyme is believed to be ubiquinone. |
| **NDUFA6** | **9606.ENSP00000482543** | NADH dehydrogenase [ubiquinone] 1 alpha subcomplex subunit 6; Accessory subunit of the mitochondrial membrane respiratory chain NADH dehydrogenase (Complex I), that is believed to be not involved in catalysis. Required for proper complex I assembly. Complex I functions in the transfer of electrons from NADH to the respiratory chain. The immediate electron acceptor for the enzyme is believed to be ubiquinone. |
| **NDUFA8** | **9606.ENSP00000362873** | NADH dehydrogenase [ubiquinone] 1 alpha subcomplex subunit 8; Accessory subunit of the mitochondrial membrane respiratory chain NADH dehydrogenase (Complex I), that is believed not to be involved in catalysis. Complex I functions in the transfer of electrons from NADH to the respiratory chain. The immediate electron acceptor for the enzyme is believed to be ubiquinone. |
| **NDUFA9** | **9606.ENSP00000266544** | NADH dehydrogenase [ubiquinone] 1 alpha subcomplex subunit 9, mitochondrial; Accessory subunit of the mitochondrial membrane respiratory chain NADH dehydrogenase (Complex I), that is believed not to be involved in catalysis. Required for proper complex I assembly. Complex I functions in the transfer of electrons from NADH to the respiratory chain. The immediate electron acceptor for the enzyme is believed to be ubiquinone. |
| **NDUFB1** | **9606.ENSP00000483888** | NADH dehydrogenase [ubiquinone] 1 beta subcomplex subunit 1; Accessory subunit of the mitochondrial membrane respiratory chain NADH dehydrogenase (Complex I) that is believed not to be involved in catalysis. Complex I functions in the transfer of electrons from NADH to the respiratory chain. The immediate electron acceptor for the enzyme is believed to be ubiquinone. |
| **NDUFB2** | **9606.ENSP00000419087** | NADH dehydrogenase [ubiquinone] 1 beta subcomplex subunit 2, mitochondrial; Accessory subunit of the mitochondrial membrane respiratory chain NADH dehydrogenase (Complex I), that is believed not to be involved in catalysis. Complex I functions in the transfer of electrons from NADH to the respiratory chain. The immediate electron acceptor for the enzyme is believed to be ubiquinone. |

| | | |
|---|---|---|
| **NDUFB3** | **9606.ENSP00000407336** | NADH dehydrogenase [ubiquinone] 1 beta subcomplex subunit 3; Accessory subunit of the mitochondrial membrane respiratory chain NADH dehydrogenase (Complex I), that is believed not to be involved in catalysis. Complex I functions in the transfer of electrons from NADH to the respiratory chain. The immediate electron acceptor for the enzyme is believed to be ubiquinone. |
| **NDUFB4** | **9606.ENSP00000184266** | NADH dehydrogenase [ubiquinone] 1 beta subcomplex subunit 4; Accessory subunit of the mitochondrial membrane respiratory chain NADH dehydrogenase (Complex I), that is believed not to be involved in catalysis. Complex I functions in the transfer of electrons from NADH to the respiratory chain. The immediate electron acceptor for the enzyme is believed to be ubiquinone. |
| **NDUFB6** | **9606.ENSP00000369176** | NADH dehydrogenase [ubiquinone] 1 beta subcomplex subunit 6; Accessory subunit of the mitochondrial membrane respiratory chain NADH dehydrogenase (Complex I), that is believed not to be involved in catalysis. Complex I functions in the transfer of electrons from NADH to the respiratory chain. The immediate electron acceptor for the enzyme is believed to be ubiquinone. |
| **NDUFB7** | **9606.ENSP00000215565** | NADH dehydrogenase [ubiquinone] 1 beta subcomplex subunit 7; Accessory subunit of the mitochondrial membrane respiratory chain NADH dehydrogenase (Complex I), that is believed not to be involved in catalysis. Complex I functions in the transfer of electrons from NADH to the respiratory chain. The immediate electron acceptor for the enzyme is believed to be ubiquinone. |
| **NDUFS4** | **9606.ENSP00000296684** | NADH dehydrogenase [ubiquinone] iron-sulfur protein 4, mitochondrial; Accessory subunit of the mitochondrial membrane respiratory chain NADH dehydrogenase (Complex I), that is believed not to be involved in catalysis. Complex I functions in the transfer of electrons from NADH to the respiratory chain. The immediate electron acceptor for the enzyme is believed to be ubiquinone. |
| **NDUFS6** | **9606.ENSP00000274137** | NADH dehydrogenase [ubiquinone] iron-sulfur protein 6, mitochondrial; Accessory subunit of the mitochondrial membrane respiratory chain NADH dehydrogenase (Complex I), that is believed not to be involved in catalysis. Complex I functions in the transfer of electrons from NADH to the respiratory chain. The immediate electron acceptor for the enzyme is believed to be ubiquinone. |
| **NDUFS7** | **9606.ENSP00000233627** | NADH dehydrogenase [ubiquinone] iron-sulfur protein 7, mitochondrial; Core subunit of the mitochondrial membrane respiratory chain NADH dehydrogenase (Complex I) that is believed to belong to the minimal assembly required for catalysis. Complex I functions in the transfer of electrons from NADH to the respiratory chain. The immediate electron acceptor for the enzyme is believed to be ubiquinone. |
| **NDUFV2** | **9606.ENSP00000327268** | NADH dehydrogenase [ubiquinone] flavoprotein 2, mitochondrial; Core subunit of the mitochondrial membrane respiratory chain NADH dehydrogenase (Complex I) that is believed to belong to the minimal assembly required for catalysis. Complex I functions in the transfer of electrons from NADH to the respiratory chain. The immediate electron acceptor for the enzyme is believed to be ubiquinone (By similarity). |
| **NLN** | **9606.ENSP00000370372** | Neurolysin, mitochondrial; Hydrolyzes oligopeptides such as neurotensin, bradykinin and dynorphin A; Belongs to the peptidase M3 family. |
| **OTC** | **9606.ENSP00000039007** | Ornithine carbamoyltransferase, mitochondrial; Ornithine carbamoyltransferase; Belongs to the aspartate/ornithine carbamoyltransferase superfamily. OTCase family. |
| **PC** | **9606.ENSP00000498994** | Pyruvate carboxylase, mitochondrial; Pyruvate carboxylase catalyzes a 2-step reaction, involving the ATP-dependent carboxylation of the covalently attached biotin in the first step and the transfer of the carboxyl group to pyruvate in the second. Catalyzes in a tissue specific manner, the initial reactions of glucose (liver, kidney) and lipid (adipose tissue, liver, brain) synthesis from pyruvate. |
| **PDE12** | **9606.ENSP00000309142** | 2',5'-phosphodiesterase 12; Enzyme that cleaves 2',5'-phosphodiester bond linking adenosines of the 5'-triphosphorylated oligoadenylates, triphosphorylated oligoadenylates referred as 2-5A modulates the 2-5A system. Degrades triphosphorylated 2-5A to produce AMP and ATP. Also cleaves 3',5'-phosphodiester bond of oligoadenylates. Plays a role as a negative regulator of the 2-5A system that is one of the major pathways for antiviral and antitumor functions induced by interferons (IFNs). |
| **PGS1** | **9606.ENSP00000262764** | CDP-diacylglycerol--glycerol-3-phosphate 3-phosphatidyltransferase, mitochondrial; Functions in the biosynthesis of the anionic phospholipids phosphatidylglycerol and cardiolipin; Belongs to the CDP-alcohol phosphatidyltransferase class-II family. |
| **PITRM1** | **9606.ENSP00000370377** | Presequence protease, mitochondrial; Metalloendopeptidase of the mitochondrial matrix that functions in peptide cleavage and degradation rather than in protein processing. Has an ATP-independent activity. Specifically cleaves peptides in the range of 5 to 65 residues. Shows a preference for cleavage after small polar residues and before basic residues, but without any positional preference. Degrades the transit peptides of mitochondrial proteins after their cleavage. Also degrades other unstructured peptides. |
| **PMAIP1** | **9606.ENSP00000326119** | Phorbol-12-myristate-13-acetate-induced protein 1; Promotes activation of caspases and apoptosis. Promotes mitochondrial membrane changes and efflux of apoptogenic proteins from the mitochondria. Contributes to p53/TP53-dependent apoptosis after radiation exposure. Promotes proteasomal degradation of MCL1. Competes with BAK1 for binding to MCL1 and can displace BAK1 from its binding site on MCL1 (By similarity). Competes with BIM/BCL2L11 for binding to MCL1 and can displace BIM/BCL2L11 from its binding site on MCL1. Belongs to the PMAIP1 family. |
| **POLG2** | **9606.ENSP00000442563** | DNA polymerase subunit gamma-2, mitochondrial; Mitochondrial polymerase processivity subunit. Stimulates the polymerase and exonuclease activities, and increases the processivity of the enzyme. Binds to ss-DNA. |

| | | |
|---|---|---|
| **POLRMT** | **9606.ENSP00000465759** | DNA-directed RNA polymerase, mitochondrial; DNA-dependent RNA polymerase catalyzes the transcription of mitochondrial DNA into RNA using the four ribonucleoside triphosphates as substrates. Component of the mitochondrial transcription initiation complex, composed at least of TFB2M, TFAM and POLRMT that is required for basal transcription of mitochondrial DNA. In this complex, TFAM recruits POLRMT to a specific promoter whereas TFB2M induces structural changes in POLRMT to enable promoter opening and trapping of the DNA non-template strand. |
| **PYCR1** | **9606.ENSP00000384949** | Pyrroline-5-carboxylate reductase 1, mitochondrial; Housekeeping enzyme that catalyzes the last step in proline biosynthesis. Can utilize both NAD and NADP, but has higher affinity for NAD. Involved in the cellular response to oxidative stress. Belongs to the pyrroline-5-carboxylate reductase family. |
| **RMND1** | **9606.ENSP00000412708** | Required for meiotic nuclear division protein 1 homolog; Required for mitochondrial translation, possibly by coordinating the assembly or maintenance of the mitochondrial ribosome; Belongs to the RMD1/sif2 family. |
| **SCO1** | **9606.ENSP00000255390** | Protein SCO1 homolog, mitochondrial; Copper metallochaperone essential for the maturation of cytochrome c oxidase subunit II (MT-CO2/COX2). Not required for the synthesis of MT-CO2/COX2 but plays a crucial role in stabilizing MT- CO2/COX2 during its subsequent maturation. Involved in transporting copper to the Cu(A) site on MT-CO2/COX2. Plays an important role in the regulation of copper homeostasis by controlling the abundance and cell membrane localization of copper transporter CTR1 (By similarity). Belongs to the SCO1/2 family. |
| **SDHA** | **9606.ENSP00000264932** | Succinate dehydrogenase [ubiquinone] flavoprotein subunit, mitochondrial; Flavoprotein (FP) subunit of succinate dehydrogenase (SDH) that is involved in complex II of the mitochondrial electron transport chain and is responsible for transferring electrons from succinate to ubiquinone (coenzyme Q). Can act as a tumor suppressor; Belongs to the FAD-dependent oxidoreductase 2 family. FRD/SDH subfamily. |
| **SELENOO** | **9606.ENSP00000370288** | Protein adenylyltransferase SelO, mitochondrial; Catalyzes the transfer of adenosine 5'-monophosphate (AMP) to Ser, Thr and Tyr residues of target proteins (AMPylation). May be a redox-active mitochondrial selenoprotein which interacts with a redox target protein. Belongs to the SELO family. |
| **SLC25A11** | **9606.ENSP00000225665** | Mitochondrial 2-oxoglutarate/malate carrier protein; Catalyzes the transport of 2-oxoglutarate across the inner mitochondrial membrane in an electroneutral exchange for malate or other dicarboxylic acids, and plays an important role in several metabolic processes, including the malate-aspartate shuttle, the oxoglutarate/isocitrate shuttle, in gluconeogenesis from lactate, and in nitrogen metabolism. Maintains mitochondrial fusion and fission events, and the organization and morphology of cristae. Involved in the regulation of apoptosis (By similarity). Acts as a tumor-suppressor gene. |
| **SLC25A13** | **9606.ENSP00000400101** | Calcium-binding mitochondrial carrier protein Aralar2; Mitochondrial and calcium-binding carrier that catalyzes the calcium-dependent exchange of cytoplasmic glutamate with mitochondrial aspartate across the mitochondrial inner membrane. May have a function in the urea cycle. |
| **SSBP1** | **9606.ENSP00000419665** | Single-stranded DNA-binding protein, mitochondrial; Binds preferentially and cooperatively to pyrimidine rich single-stranded DNA (ss-DNA). In vitro, required to maintain the copy number of mitochondrial DNA (mtDNA) and plays crucial roles during mtDNA replication that stimulate activity of the replisome components POLG and TWNK at the replication fork. Promotes the activity of the gamma complex polymerase POLG, largely by organizing the template DNA and eliminating secondary structures to favor ss-DNA conformations that facilitate POLG activity. |
| **SURF1** | **9606.ENSP00000361042** | Surfeit locus protein 1; Component of the MITRAC (mitochondrial translation regulation assembly intermediate of cytochrome c oxidase complex) complex, that regulates cytochrome c oxidase assembly. |
| **TAMM41** | **9606.ENSP00000398596** | Phosphatidate cytidylyltransferase, mitochondrial; Catalyzes the formation of CDP-diacylglycerol (CDP-DAG) from phosphatidic acid (PA) in the mitochondrial inner membrane. Required for the biosynthesis of the dimeric phospholipid cardiolipin, which stabilizes supercomplexes of the mitochondrial respiratory chain in the mitochondrial inner membrane. |
| **TEFM** | **9606.ENSP00000462963** | Transcription elongation factor, mitochondrial; Transcription elongation factor which increases mitochondrial RNA polymerase processivity. Regulates transcription of the mitochondrial genome, including genes important for the oxidative phosphorylation machinery; Belongs to the TEFM family. |
| **TFAM** | **9606.ENSP00000420588** | Transcription factor A, mitochondrial; Binds to the mitochondrial light strand promoter and functions in mitochondrial transcription regulation. Component of the mitochondrial transcription initiation complex, composed at least of TFB2M, TFAM and POLRMT that is required for basal transcription of mitochondrial DNA. In this complex, TFAM recruits POLRMT to a specific promoter whereas TFB2M induces structural changes in POLRMT to enable promoter opening and trapping of the DNA non-template strand. Required for accurate and efficient promoter recognition by the mitochondrial RNA polymerase. |
| **TFB2M** | **9606.ENSP00000355471** | Dimethyladenosine transferase 2, mitochondrial; S-adenosyl-L-methionine-dependent rRNA methyltransferase which may methylate two specific adjacent adenosines in the loop of a conserved hairpin near the 3'-end of 12S mitochondrial rRNA (Probable). Component of the mitochondrial transcription initiation complex, composed at least of TFB2M, TFAM and POLRMT that is required for basal transcription of mitochondrial DNA. In this complex, TFAM recruits POLRMT to a specific promoter whereas TFB2M induces structural changes in POLRMT to enable promoter opening and trapping of the DNA non- template. |
| **TMEM177** | **9606.ENSP00000402661** | Transmembrane protein 177; Plays a role in the early steps of cytochrome c oxidase subunit II (MT-CO2/COX2) maturation and is required for the stabilization of COX20 and the newly synthesized MT-CO2/COX2 protein. |

| | | |
|---|---|---|
| **TOP3A** | **9606.ENSP00000321636** | DNA topoisomerase 3-alpha; Releases the supercoiling and torsional tension of DNA introduced during the DNA replication and transcription by transiently cleaving and rejoining one strand of the DNA duplex. Introduces a single-strand break via transesterification at a target site in duplex DNA. The scissile phosphodiester is attacked by the catalytic tyrosine of the enzyme, resulting in the formation of a DNA-(5'-phosphotyrosyl)- enzyme intermediate and the expulsion of a 3'-OH DNA strand. The free DNA strand then undergoes passage around the unbroken strand thus removing DNA supercoils. |
| **TWNK** | **9606.ENSP00000309595** | Twinkle protein, mitochondrial; Involved in mitochondrial DNA (mtDNA) metabolism. Could function as an adenine nucleotide-dependent DNA helicase. Function inferred to be critical for lifetime maintenance of mtDNA integrity. In vitro, forms in combination with POLG, a processive replication machinery, which can use double-stranded DNA (dsDNA) as template to synthesize single-stranded DNA (ssDNA) molecules. May be a key regulator of mtDNA copy number in mammals. |
| **TXNRD1** | **9606.ENSP00000434516** | Thioredoxin reductase 1, cytoplasmic; Isoform 1 may possess glutaredoxin activity as well as thioredoxin reductase activity and induces actin and tubulin polymerization, leading to formation of cell membrane protrusions. Isoform 4 enhances the transcriptional activity of estrogen receptors alpha and beta while isoform 5 enhances the transcriptional activity of the beta receptor only. Isoform 5 also mediates cell death induced by a combination of interferon-beta and retinoic acid. |
| **UQCR10** | **9606.ENSP00000332887** | Cytochrome b-c1 complex subunit 9; Component of the ubiquinol-cytochrome c oxidoreductase, a multisubunit transmembrane complex that is part of the mitochondrial electron transport chain which drives oxidative phosphorylation. The respiratory chain contains 3 multisubunit complexes succinate dehydrogenase (complex II, CII), ubiquinol-cytochrome c oxidoreductase (cytochrome b-c1 complex, complex III, CIII) and cytochrome c oxidase (complex IV, CIV), that cooperate to transfer electrons derived from NADH and succinate to molecular oxygen. |
| **UQCRB** | **9606.ENSP00000430494** | Cytochrome b-c1 complex subunit 7; Component of the ubiquinol-cytochrome c oxidoreductase, a multisubunit transmembrane complex that is part of the mitochondrial electron transport chain which drives oxidative phosphorylation. The respiratory chain contains 3 multisubunit complexes succinate dehydrogenase (complex II, CII), ubiquinol-cytochrome c oxidoreductase (cytochrome b-c1 complex, complex III, CIII) and cytochrome c oxidase (complex IV, CIV), that cooperate to transfer electrons derived from NADH and succinate to molecular oxygen. |
| **UQCRFS1** | **9606.ENSP00000306397** | Cytochrome b-c1 complex subunit Rieske, mitochondrial; [Cytochrome b-c1 complex subunit Rieske, mitochondrial]: Component of the ubiquinol-cytochrome c oxidoreductase, a multisubunit transmembrane complex that is part of the mitochondrial electron transport chain which drives oxidative phosphorylation. The respiratory chain contains 3 multisubunit complexes succinate dehydrogenase (complex II, CII), ubiquinol-cytochrome c oxidoreductase (cytochrome b- c1 complex, complex III, CIII) and cytochrome c oxidase (complex IV, CIV), that cooperate to transfer electrons derived from NADH and succinate to molecular oxygen. |
